# Systematic genetic dissection of chitin degradation and uptake in *Vibrio cholerae*

**DOI:** 10.1101/137042

**Authors:** Chelsea A. Hayes, Triana N. Dalia, Ankur B. Dalia

**Affiliations:** Department of Biology, Indiana University, Bloomington, Indiana, USA

## Abstract

*Vibrio cholerae* is a natural resident of the aquatic environment, where a common nutrient is the chitinous exoskeletons of microscopic crustaceans. Chitin utilization requires chitinases, which degrade this insoluble polymer into soluble chitin oligosaccharides. These oligosaccharides also serve as an inducing cue for natural transformation in *Vibrio* species. There are 7 predicted endochitinase-like genes in the *V. cholerae* genome. Here, we systematically dissect the contribution of each gene to growth on chitin as well as induction of natural transformation. Specifically, we created a strain that lacks all 7 putative chitinases and from this strain, generated a panel of strains where each expresses a single chitinase. We also generated expression plasmids to ectopically express all 7 chitinases in our chitinase deficient strain. Through this analysis, we found that low levels of chitinase activity are sufficient for natural transformation, while growth on insoluble chitin as a sole carbon source requires more robust and concerted chitinase activity. We also assessed the role that the three uptake systems for the chitin degradation products GlcNAc, (GlcNAc)_2_, and (GlcN)_2_, play in chitin utilization and competence induction. Cumulatively, this study provides mechanistic details for how this pathogen utilizes chitin to thrive and evolve in its environmental reservoir.

**ORIGINALITY-SIGNIFICANCE STATEMENT:** *Vibrio cholerae*, the causative agent of the diarrheal disease cholera, interacts with the chitinous shells of crustacean zooplankton in the aquatic environment, which serves as an environmental reservoir for this pathogen. It degrades and utilizes chitin-derived products as a source of carbon and nitrogen. Also, chitin serves as an inducing cue for natural transformation – an important mechanism of horizontal gene transfer in this species. Here, we systematically dissect the genes required for chitin degradation and uptake, and characterize the role of these genes for growth on chitin as a nutrient and during chitin-induced natural transformation. Thus, this study provides mechanistic details for how this pathogen utilizes chitin to thrive and evolve in its environmental reservoir.

## INTRODUCTION

The cholera pathogen, *Vibrio cholerae*, is a natural resident of the aquatic environment. In this niche, this bacterium forms biofilms on the chitinous shells of crustacean zooplankton. These chitin biofilms are important for the water-borne transmission of cholera (Colwell et al., 2003). Also, chitin, an insoluble polymer of β1,4-linked N-acetylglucosamine (GlcNAc), serves as an important carbon and nitrogen source for *V. cholerae* in the environment (Huq et al., 1983). To utilize this carbon source, this pathogen must degrade chitin into soluble oligosaccharides via the action of chitinases. Subsequently, these chitin oligosaccharides are transported across the outer membrane and into the periplasm via a chitoporin and further broken down into mono- and di-saccharides, which can be transported across the inner membrane by specific transporters (Meibom et al., 2004; Hunt et al., 2008).

Chitin oligosaccharides also induce the genes required for natural transformation in *V. cholerae*, a physiological state in which cells can take up exogenous DNA and integrate it into their chromosome by homologous recombination (Meibom et al., 2005). Therefore, the interaction of *V. cholerae* with chitin is important for the survival and evolution of this pathogen in its environmental reservoir as well as transmission to its human host. The chitin utilization genes of *V. cholerae* have been identified by homology as well as by identifying genes induced in the presence of chitin oligosaccharides (Meibom et al., 2004; Hunt et al., 2008). For degradation, *V. cholerae* encodes 7 putative extracellular endochitinase genes and 3 putative periplasmic exochitinases (Li and Roseman, 2004; Hunt et al., 2008). Endochitinases cleave within the polymer strand of insoluble chitin and liberate soluble oligosaccharides, while exochitinases cleave terminal mono- and disaccharides from soluble chitin oligosaccharides. Since secreted endochitinases carry out the initial steps in chitin degradation, we have initially focused our efforts to characterize the role that these enzymes plan in chitin degradation. The putative endochitinases in *V. cholerae* are ChiA1 (VC1952), ChiA2 (VCA0027), VC0769, VCA0700, VC1073, VCA0140, and GbpA (VCA0811). ChiA1 and ChiA2 have previously been implicated as the major chitinases required for chitin degradation (Meibom et al., 2004; Watve et al., 2015; Dalia, 2016). VC0769 and VC1073 are predicted endochitinases, however, their role in chitin degradation in *V. cholerae* has not formally been tested. VCA0700 is a predicted periplasmic chitodextrinase, which further degrades soluble chitin oligosaccharides (Keyhani and Roseman, 1996b). VCA0140 encodes a predicted spindolin-related protein, however, this gene also contains a predicted chitin-binding domain and was therefore included as a putative endochitinase. Finally, GbpA is a GlcNAc binding protein, however, it is also predicted to contain lytic polysaccharide monooxygenase activity (Loose et al., 2014). Chitinases have been shown to function cooperatively in other chitinolytic organisms to promote chitin degradation (Suzuki et al., 2002). However, a systematic analysis of chitinases has not been performed in *V. cholerae* to assess the possibility of synergy among these enzymes or the relative contribution of each to chitin-dependent growth and induction of natural transformation.

Here, we systematically dissect the genes required for chitin degradation and uptake via multiplex genome editing by natural transformation (MuGENT) (Dalia et al., 2014b). This analysis has uncovered the endochitinases that are necessary and sufficient for both chitin utilization and chitin-induced natural transformation in *V. cholerae*.

## RESULTS

### Single mutants reveal that ChiA2 is critical for growth on chitin and chitin-induced natural transformation

First, we assessed the role of each putative chitinase during growth on chitin as a sole carbon source and chitin-induced natural transformation in single mutant strains. We find that ChiA2 is important for both growth on chitin and chitin-induced natural transformation (Fig. 1A and 1B). This is consistent with recent reports on the importance of ChiA2 for chitin-induced natural transformation (Mondal and Chatterjee, 2016). WT levels of growth on chitin also required the periplasmic chitodextrinase VCA0700 (Fig. 1A), however, this gene was dispensable for chitin-induced natural transformation (Fig. 1B). Interestingly, while chitin-induced natural transformation was reduced in mutants lacking ChiA2, it was not as deficient as a mutant lacking *pilA*, which fails to make the pilus required for uptake of exogenous DNA (Seitz and Blokesch, 2013). Since soluble chitin oligosaccharides generated by chitinases are required to induce natural competence (Meibom et al., 2005), these data suggest that other chitinases may be able to support a low level of chitin-induced natural transformation in the absence of ChiA2.

**Fig. 1.**
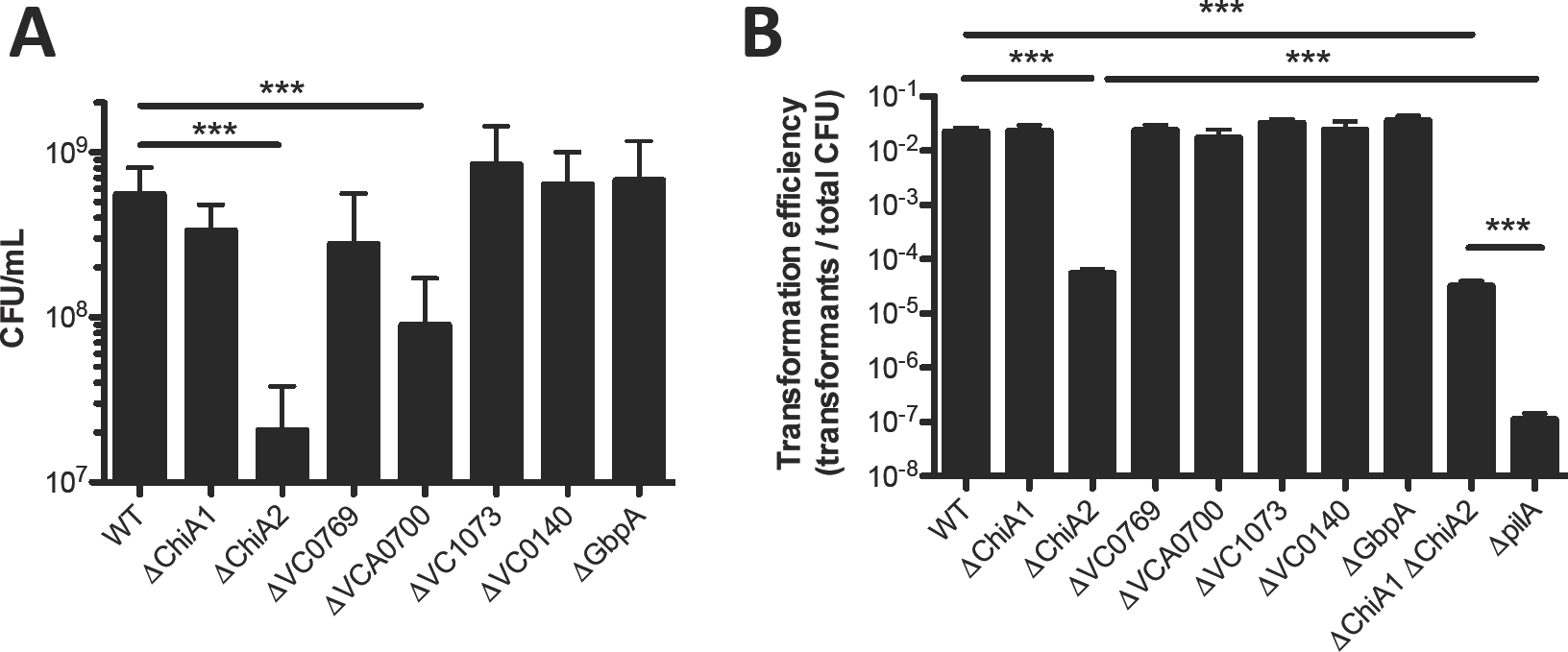
Characterizing chitinase single mutants for growth on chitin and natural transformation. (**A**) Growth of the indicated mutant strains in M9 minimal medium with chitin as a sole carbon source. (**B**) Chitin-induced natural transformation of the indicated mutant strains. All data are shown as the mean ± SD and are from at least 3 independent biological replicates. *** = *p*<0.001.

### MuGENT for systematic genetic dissection of chitinases

While our data above highlighted that ChiA2 is important for chitin degradation, we still observed chitin–dependent natural transformation in the absence of this enzyme. To determine if other endochitinases function cooperatively and/or in the absence of ChiA2, we decided to generate a strain where all 7 chitinase-like genes were inactivated. To accomplish this, we used a method we previously developed called MuGENT. This method allows for making multiple scarless mutations simultaneously in a single step (Dalia et al., 2014b; Hayes et al., 2017). The 7 chitinase-like genes are spread throughout both chromosomes (Fig. 2A), and were targeted for inactivation by generating out-of-frame ∼500bp deletions in the 5’ end of each gene. Using this approach, we rapidly generated a strain lacking all 7 putative chitinases, which we refer to as Δ7 henceforth (Fig. 2B). We then created a panel of strains where each expresses only a single chitinase (i.e. are Δ6) by systematically reverting one putative chitinase gene in each strain (Fig. 2B).

**Fig. 2.**
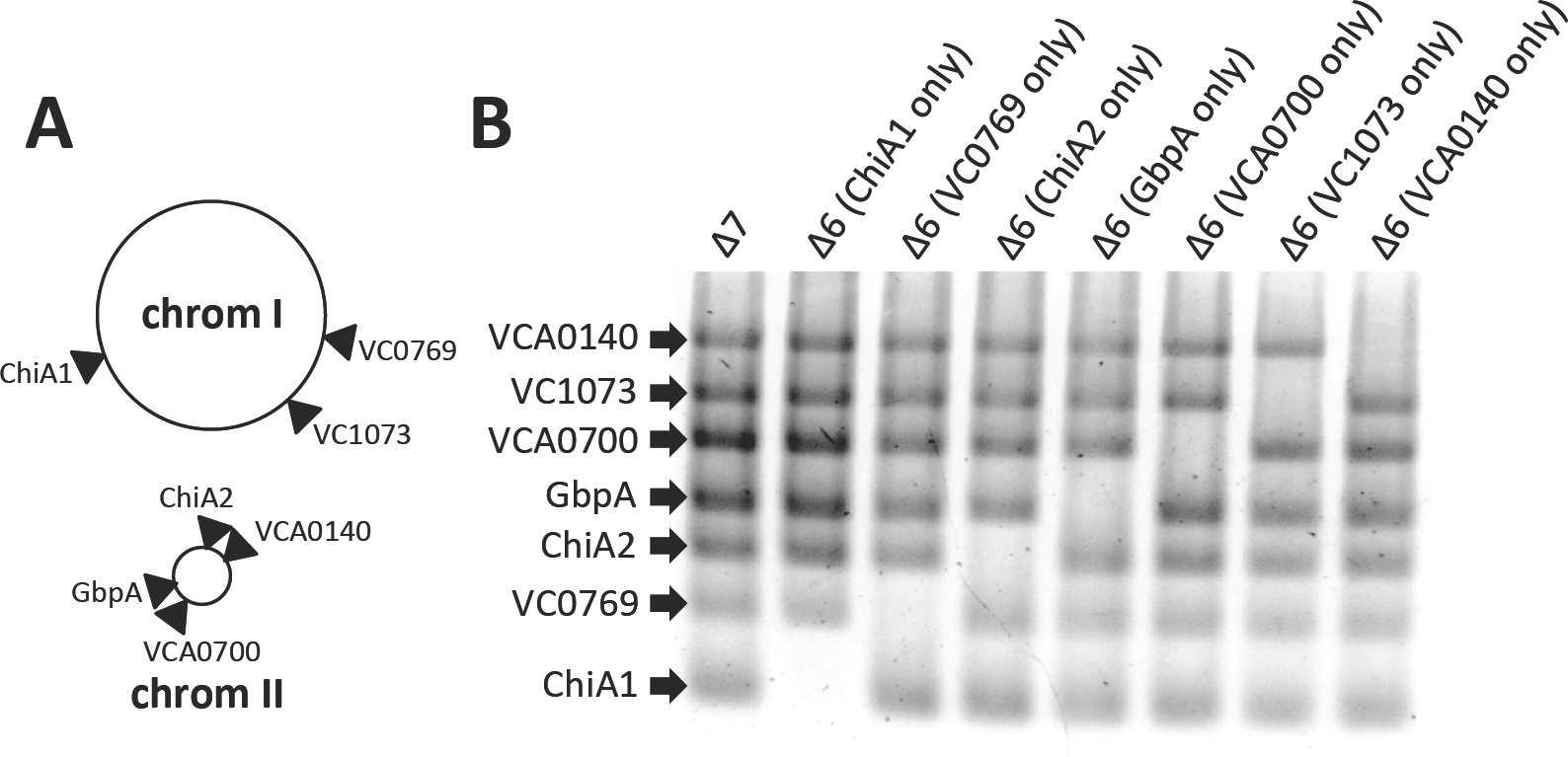
MuGENT for systematic inactivation of all 7 chitinase-like genes. (**A**) Chromosomal map of the location of the seven chitinases inactivated in this study. (**B**) MASC-PCR of the indicated mutants. The presence of a band indicates that the gene indicated to the left is inactivated, while the absence of a band indicates that this gene is intact.

### MuGENT of chitinases reveals that ChiA2 is sufficient for chitin-induced natural transformation but not growth on chitin as a sole carbon source

First, we assessed our Δ7 strain for growth on chitin and chitin-induced natural transformation. As expected, we found that this strain grew poorly on chitin (Fig. 3A). Also, we find that the Δ7 strain is significantly reduced for chitin-induced natural transformation (at the limit of detection) compared to a ChiA2 single mutant or a ChiA1 ChiA2 double mutant (Fig. 1B and 3B). This is consistent with the other chitinases playing a minor role in promoting chitin-induced natural transformation in the absence of ChiA2. Chitin induction for natural transformation can be bypassed by overexpression of TfoX, the master regulator of competence (Meibom et al., 2005). To confirm that natural transformation was only attenuated in our Δ7 strain as a result of reduced chitinase activity, we ectopically expressed TfoX in this mutant and tested natural transformation in a chitin-independent assay. Under these conditions, as expected, we found that the Δ7 strain was as transformable as the WT (Fig. 3C). Also, we would predict that the Δ7 strain would still be capable of growing on chitin degradation products. Consistent with this, we find that this strain grows as well as the wildtype on chitobiose and the chitin monomer GlcNAc (Fig. S1).

**Fig. 3.**
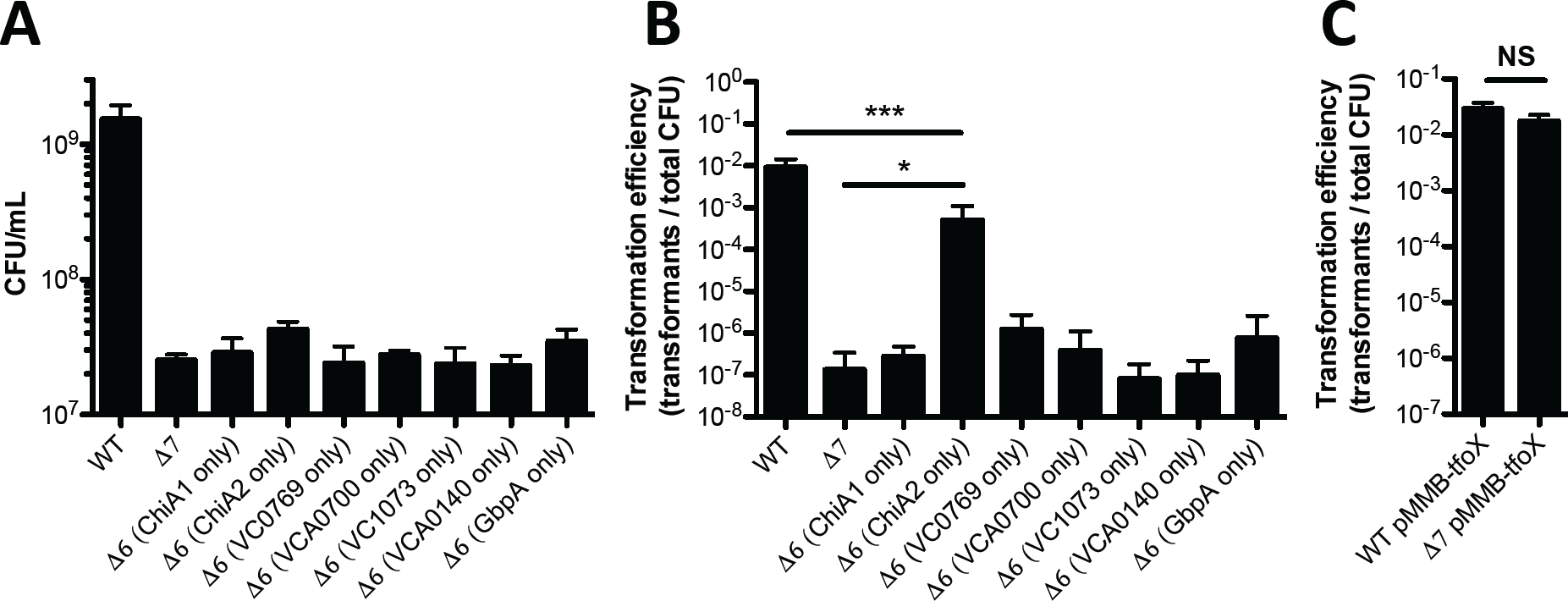
ChiA2 is sufficient for natural transformation, but not growth on chitin. (**A**) Growth of the indicated mutant strains in M9 medium with chitin as a sole carbon source. (**B**) Chitininduced natural transformation of the indicated mutant strains. (**C**) Chitin-independent natural transformation assay of the indicated mutants. TfoX was induced in these experiments with 100 μM IPTG. All data are shown as the mean ± SD and are from at least 3 independent biological replicates. * = *p*<0.05, *** = *p*<0.001, NS = not significant.

Next, we tested our panel of Δ6 mutants (each expressing a single chitinase) to determine if any chitinase was sufficient for growth on chitin and chitin-induced natural transformation. While no chitinase could independently support growth on chitin, the strain with just ChiA2 could support chitin-induced natural transformation at near WT levels (Fig. 3A and **3B**). While ChiA2 could restore chitin-induced natural transformation, none of the other chitinases could promote this activity. These results indicate that growth on chitin and chitin-induced natural transformation are separable phenomena since a strain only expressing ChiA2 supported high levels of natural transformation but did not grow on chitin as a sole carbon source. Thus, ChiA2 is both necessary and sufficient for chitininduced natural transformation.

One reason why ChiA2 may be the most important chitinase in *V. cholerae* is if this gene is simply the most highly expressed chitinase under the conditions tested. Previous work has uncovered the most highly upregulated chitinases through microarray analysis (Meibom et al., 2004), however, these studies do not provide insight into the relative expression level of each of the seven predicted endochitinases. To assess this, we performed RNA-seq on wildtype bacteria grown in the presence or absence of chitin hexasaccharide (GlcNAc)_6_ to induce the expression of chitin-regulated genes. As in prior studies (Meibom et al., 2004), lactate was provided in both conditions as a chitin-independent carbon source. Indeed, we find that ChiA2 is the most highly expressed chitinase under these conditions (Fig. 4A).

**Fig. 4.**
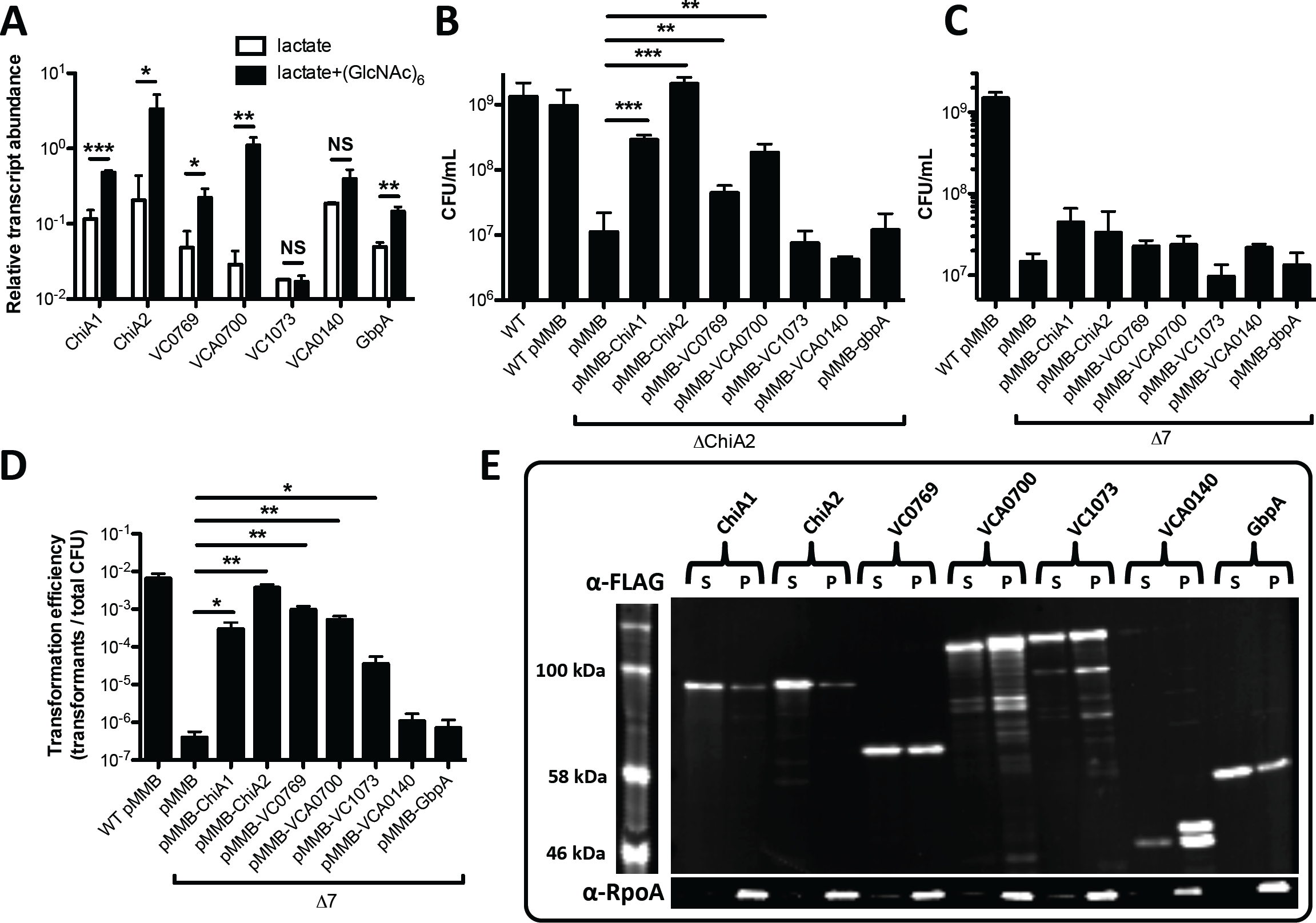
Overexpression of single chitinases in a Δ7 strain recovers natural transformation but not growth on chitin. (**A**) Relative transcript abundance of the indicated genes from RNA-seq data. (**B**) Growth of the indicated mutant strains in M9 medium containing chitin as a sole carbon source. (**C**) Growth of the indicated mutant strains in M9 medium with chitin as a sole carbon source. (**D**) Chitin-induced natural transformation of the indicated mutant strains. Genes were induced in **B**, **C** and **D** with 100 μM IPTG. (**E**) Western blot analysis of strains in the Δ7 background harboring a pMMB expression construct with the C-terminally FLAG tagged chitinase indicated. Supernatant (S) and pellet (P) fractions were run for each strain and probed with α-FLAG (top) and α-RpoA (bottom) antibodies. All data in **A**-**D** are shown as the mean ± SD and are from at least 3 independent biological replicates. * = *p* <0.05, ** = *p*<0.01, and *** = *p*<0.001, and NS = not significant.

Thus, a trivial explanation for the relative importance of ChiA2 in *V. cholerae* might be that this chitinase is the only one expressed at the levels required for efficient liberation of chitin oligosaccharides. To test this further, we bypassed the native regulation of these chitinases by ectopically expressing each in pMMB67EH (abbreviated pMMB), an IPTG-inducible P*_tac_* expression vector that supports high levels of gene expression (Furste et al., 1986). As expected, ectopic expression of ChiA2 restored wildtype levels of growth on chitin to a ChiA2 single mutant (Fig. 4B). Ectopic expression of ChiA1, VC0769, and VCA0700 also restored growth on chitin to a ChiA2 single mutant; however, this was not to wildtype levels (Fig. 4B). Thus, this result suggests that ChiA2 expression levels alone cannot fully account for the importance of this chitinase during growth on chitin and chitininduced natural transformation.

Next, we ectopically expressed each chitinase in our Δ7 mutant. This analysis uncovered that none of these chitinases, even when overexpressed, could independently support growth on chitin (Fig. 4C). However, five of these genes (*chiA1*, *chiA2*, VC0769, VCA0700, and VC1073) did enhance chitin-induced natural transformation when overexpressed (Fig. 4D). Together, these results suggest that low levels of chitinase activity may be sufficient to promote chitin-induced natural transformation, while robust (and possible concerted chitinase activity) is required for efficient chitin utilization. Furthermore, since chitininduced natural transformation requires soluble chitin oligosaccharides, this analysis suggests that VC0769, VCA0700, and VC1073 possess *bona fide* chitinase activity (Fig. 4D), however, since they could not recover wildtype levels of growth on chitin to a *chiA2* single mutant (Fig. 4B), this suggests that they may have substantially lower chitinase activity than ChiA2. Conversely, VCA0140 and GbpA did not promote chitin-induced natural transformation even when overexpressed in the Δ7 background, suggesting that these genes cannot independently liberate soluble chitin oligosaccharides (Fig. 4D). To test this further, we assessed endochitinase activity in strains where each chitinase was overexpressed in the Δ7 mutant background. Endochtinase activity was determined using remazol brilliant blue (RBB) labeled chitin beads. In this assay, liberation of soluble chitin oligosaccharides is directly correlated to the release of soluble RBB dye (Gomez Ramirez et al., 2004; Dalia, 2016). We found that the 5 chitinases that could support chitin-induced natural transformation when ectopically expressed in the Δ7 strain (Fig. 4D) all had detectable endochitinase activity with the ChiA2 expressing strain displaying the highest levels of activity (Fig. S2).

To determine expression and secretion of chitinases expressed on our pMMB constructs, we generated variants of each expression vector where each chitinase had a C-terminal FLAG tag. These FLAG-tagged constructs were functional as determined by their ability to induce natural transformation in the Δ7 mutant (Fig. S3). Western blot analysis revealed that while the total level of expression for each chitinase varied (supernatant + pellet), they were all secreted at similar levels when overexpressed (supernatant) with the exception of VCA0140, which was poorly secreted into the medium (Fig. 4E). To confirm that protein in the supernatant in this experiment was the result of secretion and not cell lysis we detected the RNA polymerase alpha subunit (RpoA), which as expected, was found only in the cell pellet fraction. Thus, these results indicate that phenotypic differences observed among the distinct chitinases likely reflects differences in activity and not differences in secretion.

### ChiA2 works in conjunction with the periplasmic chitodextrinase VCA0700 to promote growth on chitin as a sole carbon source

Since ChiA2 was not independently sufficient to promote growth on chitin as a sole carbon source, we hypothesized that it may work in conjunction with another chitinase to promote robust chitin degradation and utilization. We hypothesized that the chitodextrinase VCA0700 would be a likely candidate for two reasons. One, when characterizing the phenotypes of single mutants, we found that loss of VCA0700 resulted in reduced growth on chitin. Second, VCA0700 is predicted to be found at a distinct subcellular localization and we hypothesized that the concerted action of the extracellular chitinase ChiA2 and the predicted periplasmic chitodextrinase VCA0700 might be required for efficient chitin utilization. To test this, we took the Δ6 strain that only expressed VCA0700 and systematically knocked back in each of the other 6 chitinases to generate a panel of Δ5 strains where each is expressing 2 chitinases (one being VCA0700). When testing this panel for growth on chitin, we find that the strain that contains VCA0700 and ChiA2 is capable of wildtype levels of growth on chitin (Fig. 5A). Also, this strain displays wildtype levels of chitin-induced natural transformation (Fig. 5B). To determine if VCA0700 could work in conjunction with any of the other chitinases to mediate growth on chitin, we ectopically expressed each chitinase in the Δ6 strain that only encodes VCA0700. We found that only ChiA2 could promote robust growth in this background (Fig. 5C). Cumulatively, these results indicate that ChiA2 and VCA0700 work together to efficiently degrade insoluble chitin in *V. cholerae*.

**Fig. 5.**
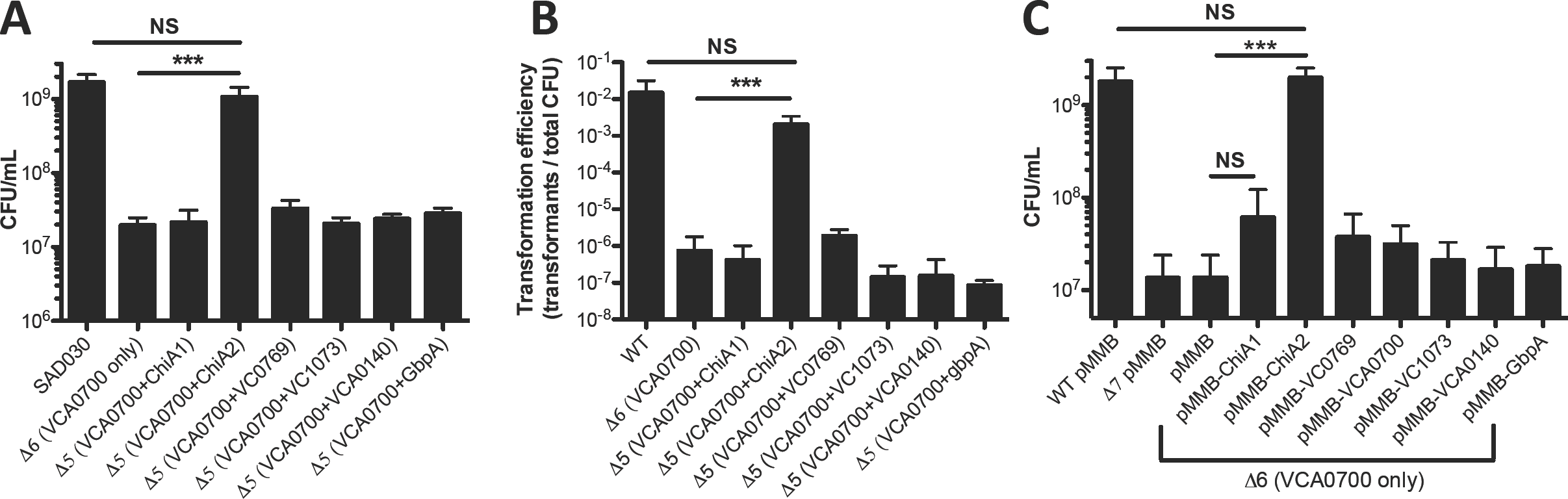
ChiA2 and VCA0700 are sufficient for growth on chitin. (**A**) Growth of the indicated mutant strains in M9 medium with chitin as a sole carbon source. (**B**) Chitin-induced natural transformation of the indicated mutant strains. (**C**) Growth of the indicated strains in M9 medium with chitin as the sole carbon source and 100 μM IPTG. All data are shown as the mean ± SD and are from at least 3 independent biological replicates. *** = *p*<0.001, NS = not significant.

### Dissecting the role of chitin transporters during growth on chitin and chitin-induced natural transformation

Once liberated from insoluble chitin via chitinase activity, chitin oligosaccharides are subsequently transported into the periplasm through the action of a chitoporin (encoded by VC0972) (Keyhani et al., 2000; Meibom et al., 2004). These oligosaccharides must then be degraded into mono- or di-saccharides that can be transported across the inner membrane. These are the monosaccharide GlcNAc and the disaccharides (GlcNAc)_2_ (i.e. chitobiose) and (GlcN)_2_ (i.e. the unacetylated chitin disaccharide). GlcNAc and (GlcN)_2_ are transported via the action of two distinct PEP-dependent phosphotransferase system transporters (VC0995 and VC1282, respectively), while (GlcNAc)_2_ is transported via an ABC transporter (permease encoded by VC0618 and VC0619) (Meibom et al., 2004; Hunt et al., 2008). To assess the role of each of these transporters during growth on chitin and chitin-induced natural transformation, we generated a panel of mutants lacking all possible combinations of the three inner membrane transporters. We also generated a strain lacking the outer membrane chitoporin. For growth on chitin, we find that the chitoporin VC0972 is required for wildtype levels of growth, which is consistent with previous reports (Fig. 6A) (Meibom et al., 2004). Also, while any one inner membrane transporter is dispensable, loss of both the GlcNAc and (GlcNAc)_2_ transporters resulted in lack of growth on chitin ( Fig. 6A). Thus, this suggests that chitin is efficiently broken down to GlcNAc and (GlcNAc)_2_, while formation of (GlcN)_2_ is less efficient. Indeed, in some sources of chitin, there is only 1 GlcN residue for every 6 GlcNAc residues (Meibom et al., 2004). While *V. cholerae* does encode a putative chitin deacetylase (VC1280) adjacent to the locus required for (GlcN)_2_ uptake, the activity or expression of this enzyme must not support robust growth on chitin as a sole carbon source (Meibom et al., 2004; Hunt et al., 2008). Among this panel of mutants, reduced growth on chitin as a sole carbon source directly correlated with reduced rates of chitin-induced natural transformation (Fig. 6A and 6B), which suggests that reduced rates of transformation among transporter mutants may largely be due to an inability to grow under competence-inducing conditions. Furthermore, we have confirmed that the defect in natural transformation among transporter mutants is specific to chitin-dependent growth and/or competence induction because all mutants were recovered for transformation by ectopic expression of TfoX (via pMMB-*tfoX*) in chitin-independent transformation assays (Fig. S4).

**Fig. 6.**
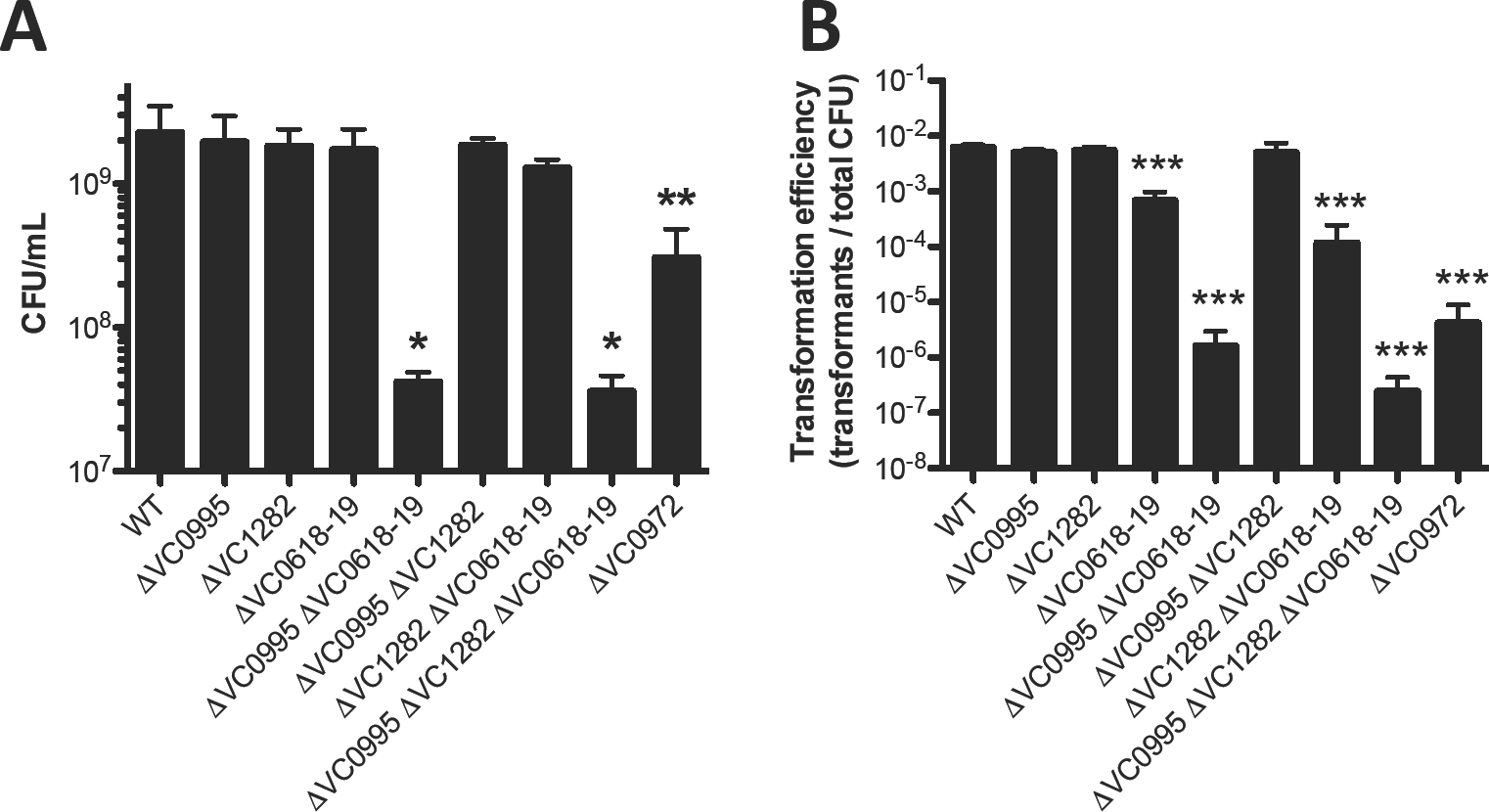
Role of chitin transporters for growth on chitin and chitin-induced natural transformation. (**A**) Growth of the indicated mutant strains in M9 medium with chitin as a sole carbon source. (**B**) Chitin-induced natural transformation of the indicated mutant strains. All data are shown as the mean ± SD and are from at least 3 independent biological replicates. All statistical comparisons in **A** and **B** were made between the indicated mutant and the WT. * = *p*<0.05, ** = *p*<0.01, and *** = *p*<0.001.

## DISCUSSION

These results suggest that ChiA2 and VCA0700 work synergistically to promote efficient degradation of chitin in *V. cholera* e for both growth on this carbon source and induction of natural transformation (Fig. 7). ChiA2 likely works extracellularly to generate chitin oligosaccharides, while the chitodextrinase VCA0700 may work in the periplasm to degrade these oligosaccharides further for uptake through the inner membrane. ChiA2 is the most important extracellular chitinase under the conditions used here, since this enzyme is necessary for growth on chitin as a sole carbon source and for chitin-induced natural transformation. Based on expression levels, ChiA2 is the most highly expressed chitinase in *V. cholerae*. Ectopic overexpression of other chitinases, however, did not complement a ChiA2 mutant, which suggests that the activity (and not just expression level) of ChiA2 may be important for the chitinolytic activity of *V. cholerae* under the conditions tested. The importance of VCA0700 for growth on chitin is consistent with previous reports, which suggested that the periplasmic steps of chitin degradation are limiting for chitin utilization (Bassler et al., 1991). Loss of VCA0700 alone, however, did not result in complete loss of growth on chitin. This suggests that in the VCA0700 mutant, the other endochitinases may work together to cleave chitin into products that can be taken up without the need for chitodextrinase activity, albeit less efficiently. Also, there are three predicted periplasmic exochitinases (VC2217, VC0613 and VC0692) that may work in concert with endochitinases to promote efficient degradation of chitin oligosaccharides in the periplasm (Fig. 7).

**Fig. 7.**
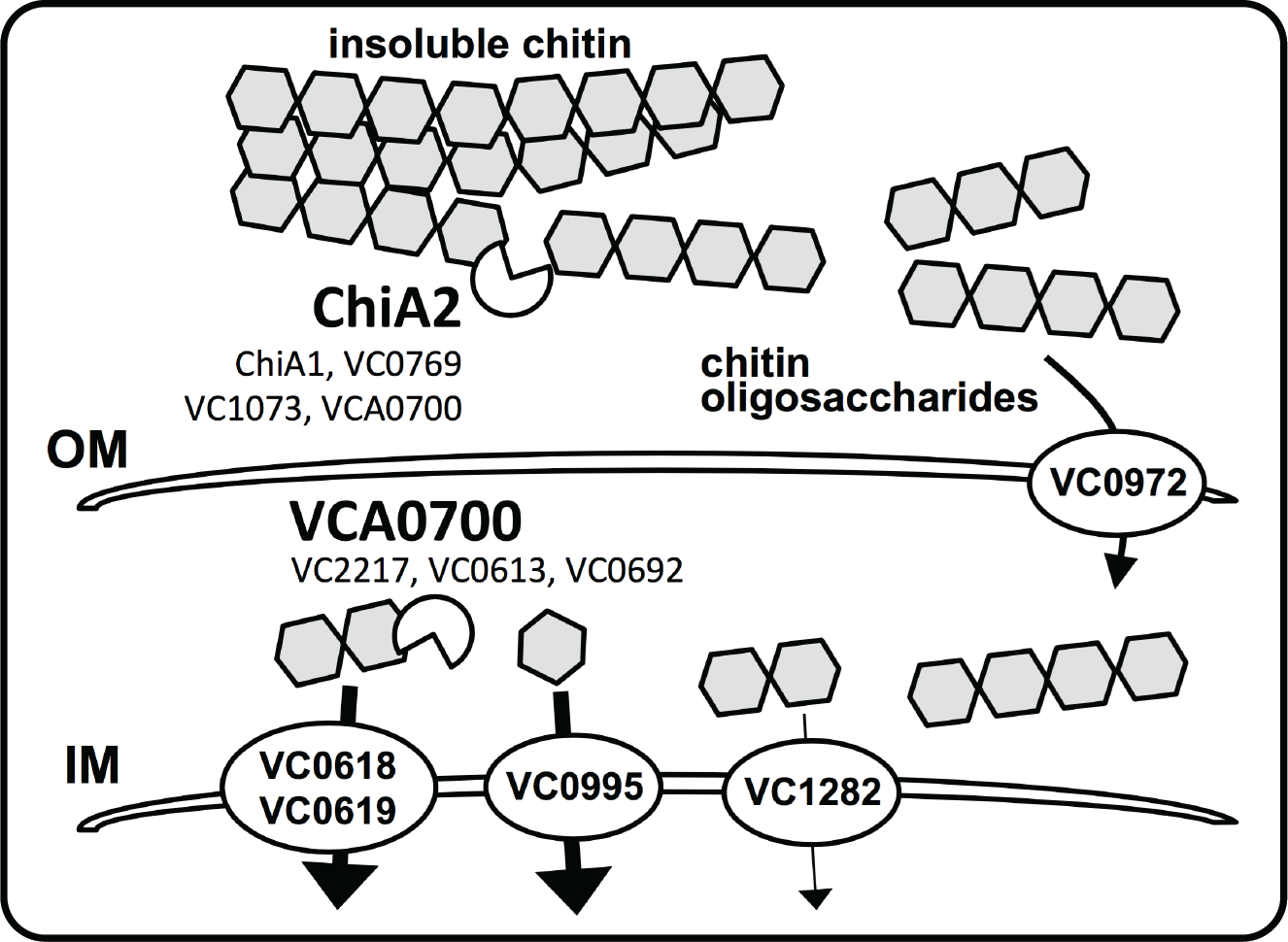
Schematic of the chitin utilization pathway genetically dissected in this study. First, extracellular chitinases degrade insoluble chitin into soluble chitin oligosaccharides. While ChiA2 is the dominant enzyme required for this process, the chitinases ChiA1, VC0769, VC1073, and VCA0700 likely play some role. These soluble oligosaccharides are then taken up across the outer membrane (OM) and into the periplasm via the chitoporin encoded by VC0972. Then, these oligosaccharides are likely further broken down by the chitodextrinase VCA0700 and/or exochitinases (VC2217, VC0613, VC0692) into (GlcNAc)_2_ (aka chitobiose), GlcNAc, and (GlcN)_2_, which are taken up across the inner membrane (IM) into the cytoplasm by the transporters encoded by VC0618-0619, VC0995, and VC1282, respectively. Our results indicate that for robust growth on chitin, the transporters responsible for uptake of chitobiose and GlcNAc play the largest role.

There are a number of reasons why degradation of chitin in two stages, one extracellular and one periplasmic, may be beneficial to the organism. One, relatively few microorganisms can take up long chitin oligosaccharides from the extracellular environment, while many microbes can take up chitin-derived mono- and di-saccharides. *Vibrio* species encode a specific chitoporin (VC0972 in *V. cholerae*) to transport oligosaccharides across the outer membrane, which could provide a competitive advantage in the environment (Suginta et al., 2013). A second benefit to this spatially segregated degradation is that chitin oligosaccharides serve as an important cue in the periplasm to signal upregulation of the chitin utilization regulon (Keyhani and Roseman, 1996a; Li and Roseman, 2004) and genes required for natural competence (Meibom et al., 2004; Dalia et al., 2014a). Thus, it is beneficial to take up long chain oligosaccharides into the periplasm to serve as an inducing cue prior to degradation for uptake and catabolism. Surprisingly, our analysis of VCA0700 indicated that when this chitodextrinase is ectopically expressed, some of this protein may be secreted to the extracellular milieu and this is not a consequence of cell lysis (Fig. 4E). Indeed, our results also indicate that VCA0700 can act extracellularly since overexpression of this chitinase supported chitin-induced natural transformation in our Δ7 strain. Thus, the concentrated action of ChiA2 and VCA0700 may be spatially segregated (extracellular ChiA2 and periplasmic VCA0700) as previously hypothesized or it is possible that both of these chitinases function extracellularly to efficiently degrade insoluble chitin into soluble oligosaccharides. Further analysis of ChiA2 and VCA0700 localization and activity will shed light on this question, which will be the focus of future work.

Induction of chitin-induced natural transformation requires ChiA2, while periplasmic degradation via the chitobextrinase VCA0700 was largely dispensable. It was previously shown that VCA0700 lacks detectable activity on insoluble chitin; however, it has robust activity on soluble long chain chitin oligosaccharides (Keyhani and Roseman, 1996b). Our previous work has shown that longer chains of chitin oligosaccharides are optimal at inducing the activity of the chitin sensor TfoS, which is required for competence induction (Meibom et al., 2005; Yamamoto et al., 2010; Dalia et al., 2014a). Thus, VCA0700 activity may not be required for competence induction since this enzyme largely acts to reduce oligosaccharide chain length while playing a limited role in liberating long chitin oligosaccharides from insoluble chitin.

Mutational analysis of the chitin transporters revealed that the outer membrane chitoporin was important for growth on chitin as a sole carbon source and for competence induction. The inner membrane GlcNAc and (GlcNAc)_2_ transporters on the other hand were genetically redundant for these activities (Fig. 7). Loss of chitin-induced natural transformation in transporter mutants directly correlated with the reduced ability of these strains to grow on chitin as a sole carbon source. A strain that only expresses the chitinase ChiA2 (i.e. a Δ6 strain), however, is not able to grow on chitin, while it displays high rates of natural transformation. Also, overexpression of 5 of the 7 predicted chitinases in the Δ7 strain supported natural transformation while none supported growth on chitin as a sole carbon source. This suggests that induction of competence only requires relatively low levels of chitinase activity and by extension only small amounts of chitin oligosaccharides, while growth on chitin may require more robust and concerted chitinase activity. As mentioned above, TfoS, the membrane-embedded chitin sensor required for natural transformation, is induced by long chain chitin oligosaccharides (Meibom et al., 2005; Yamamoto et al., 2010; Dalia et al., 2014a). Thus, loss of the chitoporin, which specifically imports long chain chitin oligosaccharides into the periplasm (Suginta et al., 2013), may result in reduced rates of natural transformation as a result of poor TfoS induction. Loss of natural transformation in the GlcNAc and (GlcNAc)_2_ transporter double mutant would not be predicted to diminish TfoS induction, however, loss of these uptake transporters may slow growth to a level where the competence machinery is no longer efficiently expressed. Alternatively, it is possible that the cytoplasmic chitin degradation products internalized by the GlcNAc and (GlcNAc)_2_ transporters aid in competence induction. However, previous work has shown that artificial activation of TfoS supports competence induction in rich medium even in the absence of chitin (Dalia et al., 2014a; Dalia, 2016).

Chitin is the second most abundant biomolecule in nature (after cellulose) and represents an abundant waste product of the seafood industry (Yan and Chen, 2015). The genes defined here may represent the minimal gene set required for efficient chitin utilization, which can be transferred to relevant non-chitinolytic microorganisms for biotech applications. This will be a focus of future work.

In conclusion, this study systematically defines the chitinases and transporters that are necessary and sufficient for chitin degradation and utilization in *V. cholerae*. Also, it identifies the unique requirements for chitin-induced horizontal gene transfer by natural transformation in this important human pathogen.

## EXPERIMENTAL PROCEDURES

### Bacterial strains and culture conditions

Strains were routinely grown in LB broth and on LB agar plates. When necessary, media was supplemented with stremptomycin (100 μg/mL), spectinomycin (200 μg/mL), kanamycin (50 μg/mL), trimethoprim (10 μg/mL), or carbenicillin (100 μg/mL). Strains in this study are all derived from E7946 (Miller et al., 1989). All strains used in this study are listed in Table S1.

For growth on chitin as the sole carbon source, we used M9 minimal medium (Difco) supplemented with 30 μM FeSO_4_ and ∼1% chitin from shrimp shells (Sigma). To 1 mL of M9+chitin medium, ∼10^5^ cells were added and grown for 48 hours at 30°C for each growth reaction. Reactions were then plated for viable counts to assess growth. For reactions with strains containing a pMMB plasmid, carbenicillin (20μg/mL) was added to M9+chitin reactions to maintain the plasmid, and IPTG (100 μM) was added to induce expression.

### Generating mutant strains and constructs

All mutants were generated by natural cotransformation and MuGENT as previously described (Dalia et al., 2014b). Briefly, mutant constructs were generated by splicing-by-overlap extension (SOE) PCR as previously described (Dalia et al., 2013). For cotransformation and MuGENT, a selected product (i.e. one conferring resistance to an antibiotic) was used as the transforming DNA in conjunction with an unselected product (i.e. one that will confer the mutation of interest). By selecting for integration of the selected product, we increase the likelihood that the unselected mutation will have integrated into cells within a competent population. We then screen for the mutation of interest in these cells by multiplex allele specific colony PCR (MASC-PCR) exactly as previously described (Wang et al., 2009). All chitinase and transporter mutations were generated using unselected SOE products. All expression constructs were generated by traditional cloning and C-terminal FLAG tags were added by site-directed mutagenesis as previously described (Edelheit et al., 2009). All primers used to generate mutant constructs and plasmids are listed in Table S2.

### Natural transformation assays

We tested chitin-induced natural transformation essentially as previously described (Dalia et al., 2015). Briefly, ∼10^8^ cells were incubated in a 1 mL reaction of instant ocean medium (7 g/L; Aquarium Systems) containing ∼8 mg of chitin. Cells were incubated in this medium at 30°C statically for 16-24 hours to induce competence. Then, transforming DNA (tDNA) was added. For all natural transformation assays in this study, we used ∼500 ng of a PCR product that would replace the frame-shifted transposase VC1807 with a trimethoprim resistance cassette. Reactions were incubated for 5-24 hours with tDNA, and then 1 mL of LB was added to outgrow reactions. The transformation efficiency was then determined by plating reactions for viable counts on LB+Tm10 (transformants) and plain LB (total viable counts).

For chitin-independent transformation assays, strains containing pMMB-tfoX were grown overnight in LB with 100 μg/mL carbenicillin and 100 μM IPTG. Then, ∼10^8^ cells were diluted into instant ocean medium containing 100 μg/mL IPTG. Next, tDNA was added and incubated statically at 30°C for 5-24 hours. Then, reactions were outgrown and plated as described above to determine the transformation efficiency.

### RNA-seq

RNA was prepped for sequencing on the Illumina platform exactly as previously described (Shishkin et al., 2015). Reads obtained were mapped to the N16961 reference genome (NC_002505 and NC_002506) and analyzed using the Tufts University Galaxy server (Afgan et al., 2016). Reads were aggregated within ORFs and normalized for the size of the ORF to obtain normalized transcript abundance for each gene under each condition tested.

### Western blot analysis

Western blots were conducted essentially as previously described (Burnette, 1981; Dalia et al., 2015). Briefly, samples were prepared for western blots by growing strains to mid-log in 20 μg/mL carbenicillin and 100μM IPTG. Cells were then spun and cell free supernatants were collected and boiled in SDS PAGE sample buffer. The cell pellets were then washed and resuspended with an equal volume of 0.5X IO and then boiled in SDS PAGE sample buffer. Samples were electrophertically separated on 10% polyacrylamide gels and transferred to a nitrocellulose membrane. FLAG-tagged proteins were probed using rabbit polyclonal α-FLAG antibodies (Sigma), while RpoA was probed using a mouse monoclonal antibody (Biolegend). Blots were developed using IRDye 800CW labeled α-rabbit or αmouse secondary antibodies as appropriate and imaged using a LI-COR Imaging system.

### Endochitinase assays

Chitin beads (New England Biolabs) were labeled with remazol brilliant blue exactly as previously described (Dalia, 2016). For each reaction, ∼10^7^ cells were added to 100uL of RBB chitin beads (50% slurry) and 600 μL of M9 minimal medium supplemented with tryptone (1%), 30 μM FeSO_4_, Carbenicillin 20 μg/mL, and 100 μM IPTG. Reactions were incubated with shaking for 72 hours at 30°C. Then, samples were centrifuged for 1 min at max speed in a microcentrifuge (21,000 × g). Next, 200 μL of the supernatant was transferred to a 96-well plate and the A_595_ was determined on a Biotek H1M plate reader.

## ACKNOWLEDGEMENTS

We would like to thank Neil Greene for critical reading of this manuscript. Also, we thank the Malcolm Winkler and Julia van Kessel labs for reagents and advice. This work was supported by US National Institutes of Health Grant AI118863 and startup funds from the Indiana University College of Arts and Sciences to ABD.

---

**Fig. S1.**
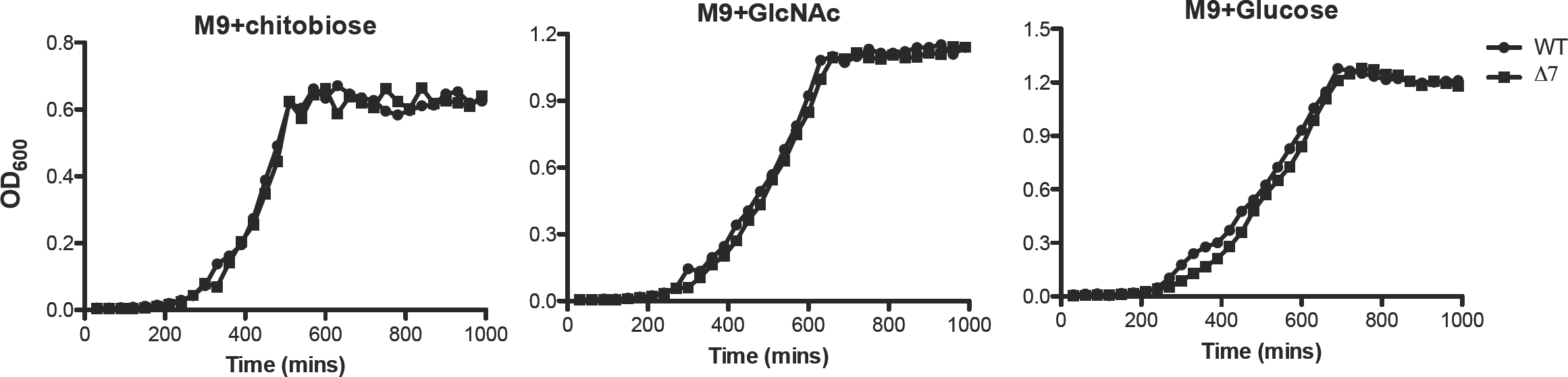
A chitinase deficient strain is still capable of growth on the chitin degradation products chitobiose and GlcNAc. Growth curves of wildtype (black circles) and Δ7 chi<nase strain (black squares) in M9 minimal medium supplemented with the carbon source indicated above each graph. Data are representa<ve of at least two independent experiments.

**Fig. S2.**
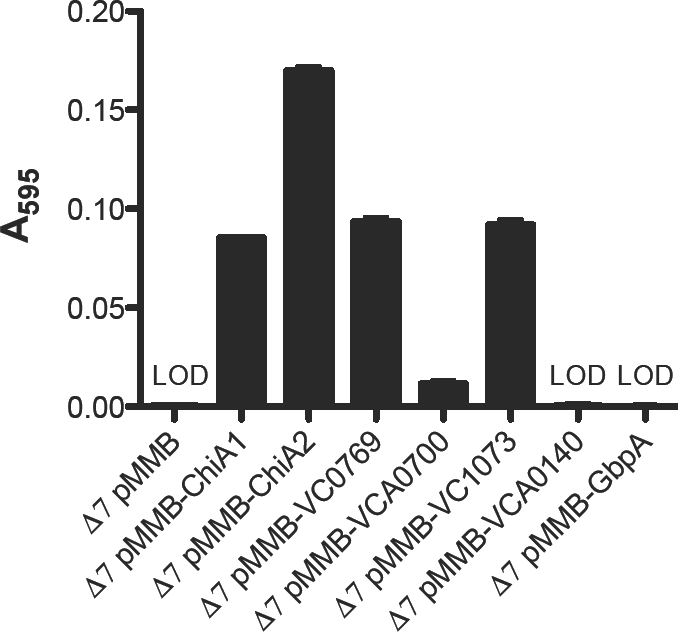
Five predicted endochitinases have detectable activity. Endochi<nase ac<vity assay of the indicated strains. All strains were incubated with RBB beads in M9+tryptone medium supplemented carbenicillin 20 μg/mL and 100 μM IPTG. LOD = limit of detec<on. Data are the result of at least three biological replicates and are shown as the mean ± SD.

**Fig. S3.**
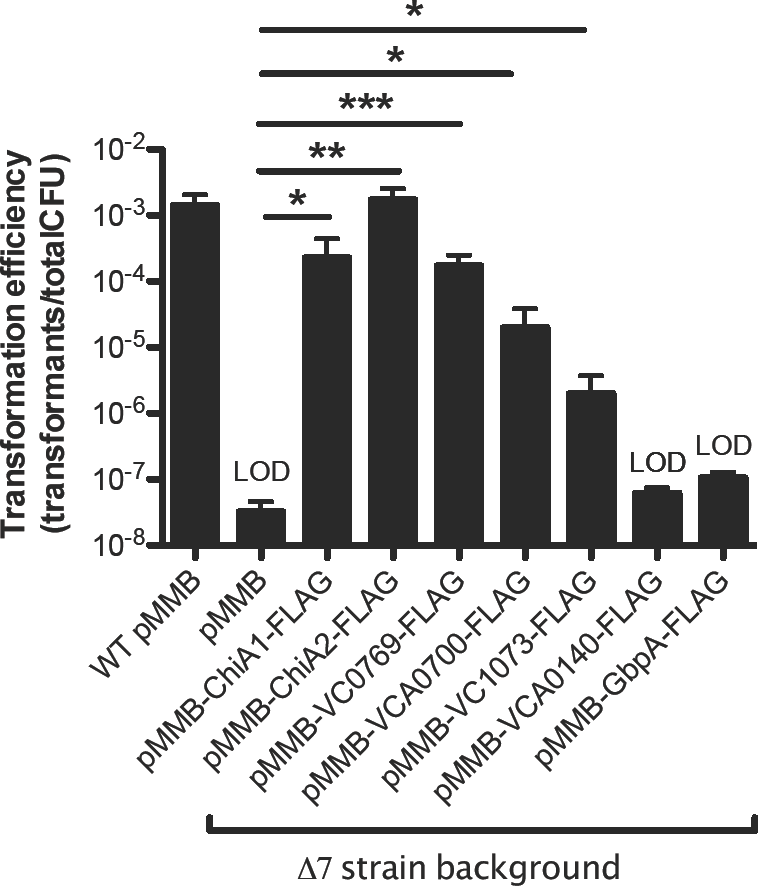
C-terminally FLAG tagged chitinases functional. Natural transforma<on assay of the indicated strains. All strains were incubated chi<n with Carbeniciillin (20 μg/mL) and IPTG(100 μg/mL). Data are from at least three independent biological replicates and shown as mean ± SD. * = p<0.05, ** = p<0.01, *** = and LOD = limit of detec<on. All data from at least three independent biological replicates and are shown as the mean ± SD.

**Fig. S4.**
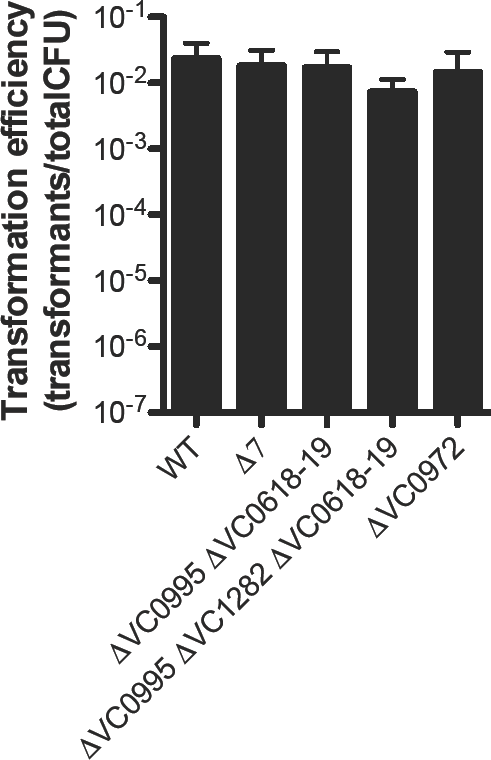
Ectopic expression of TfoX rescues transformation efficiency of transporter mutants. Chitin-independent transformation assay of the strains. All strains harbored a pMMB plasmid and were induced with 100 μM IPTG. data are from at least three independent biological replicates and are shown as the mean ± SD.

**Table S1.**
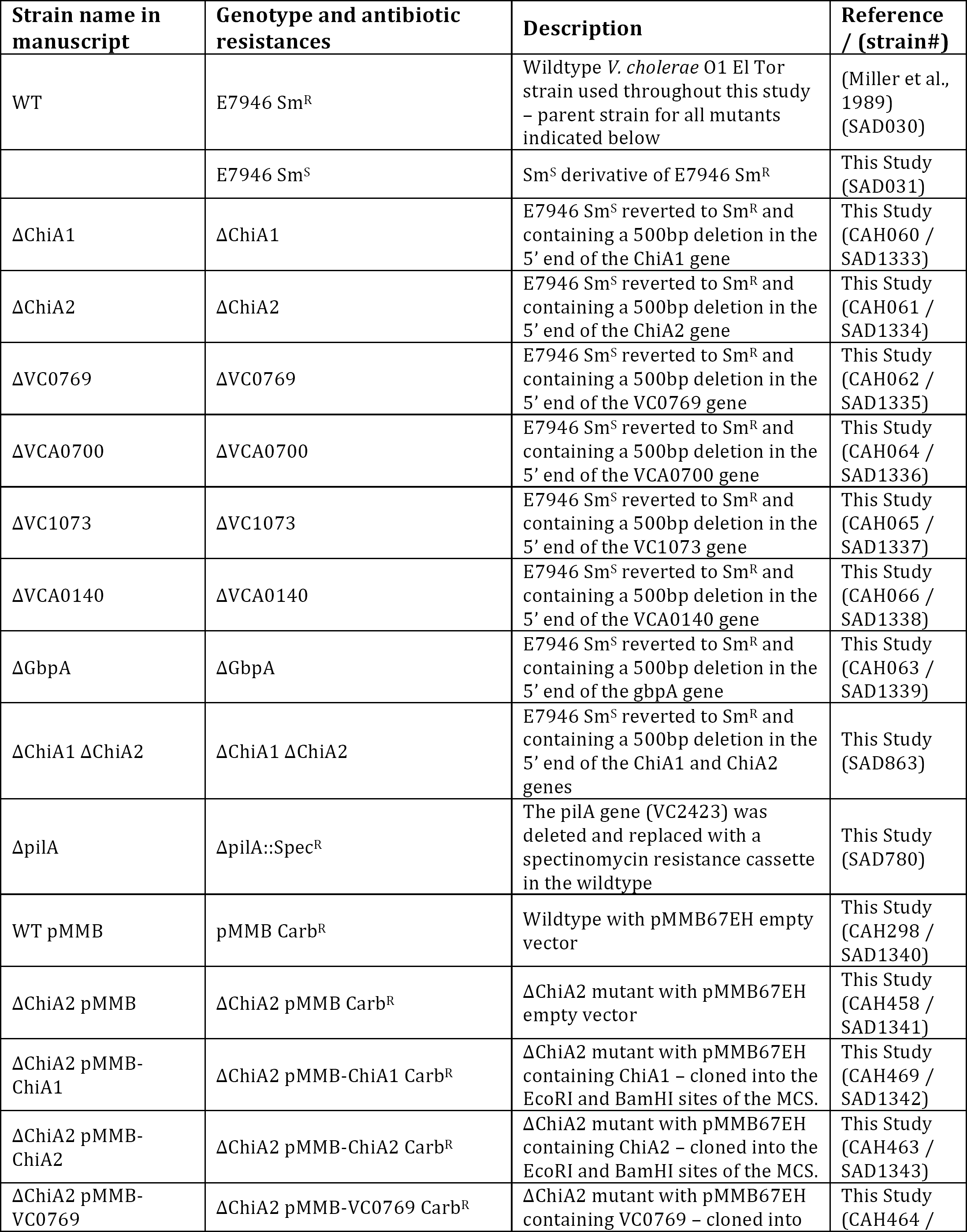
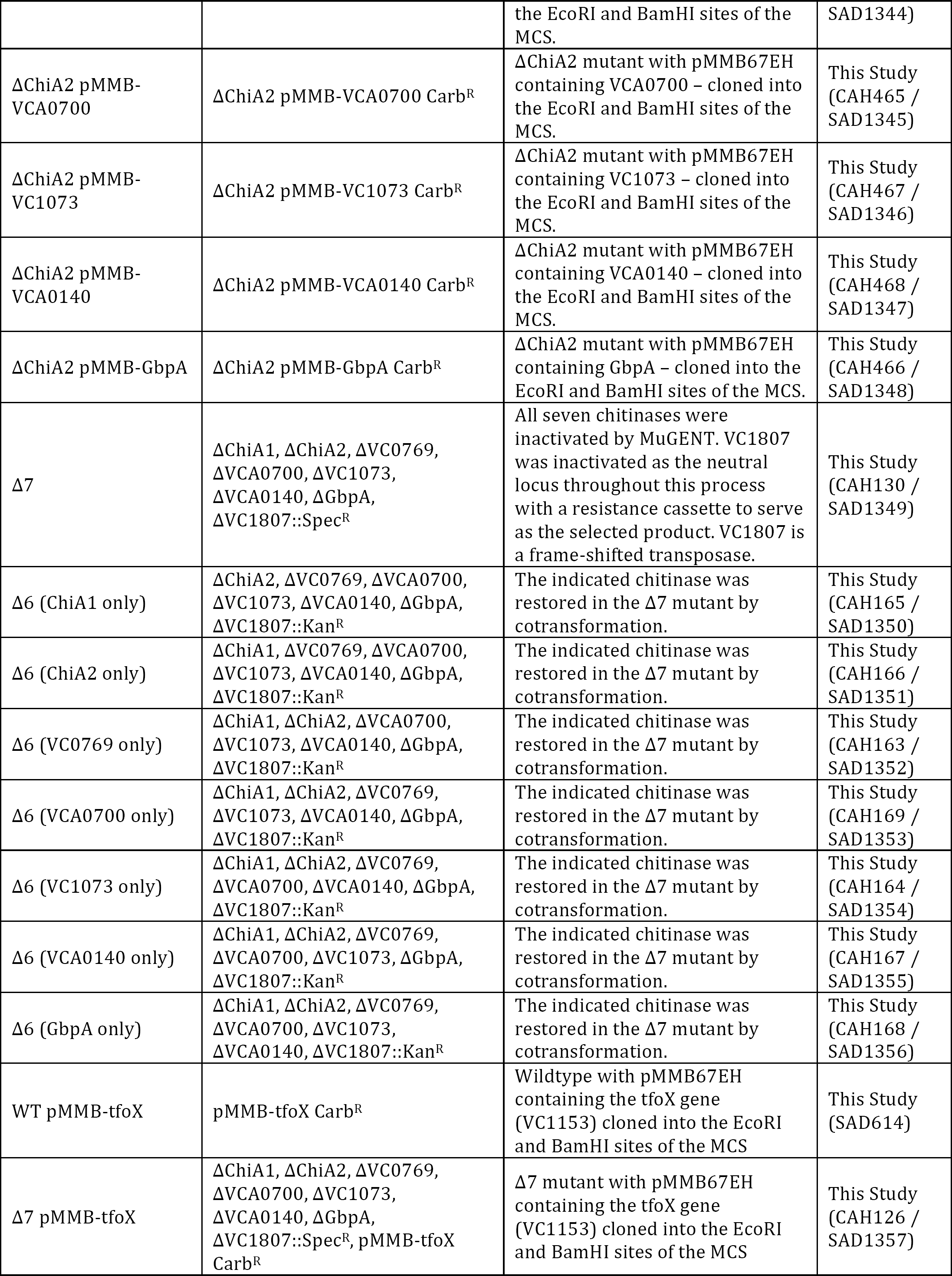
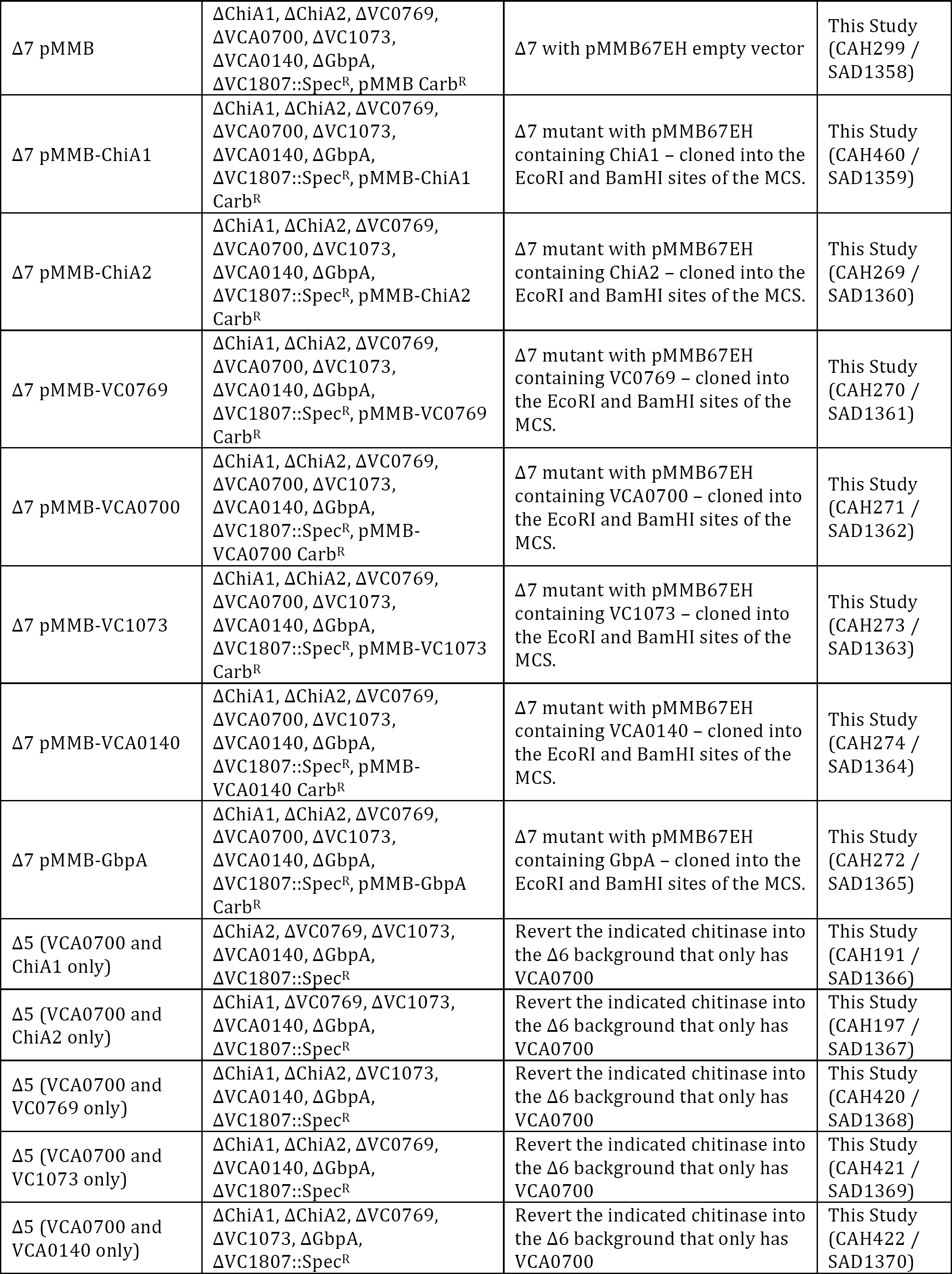
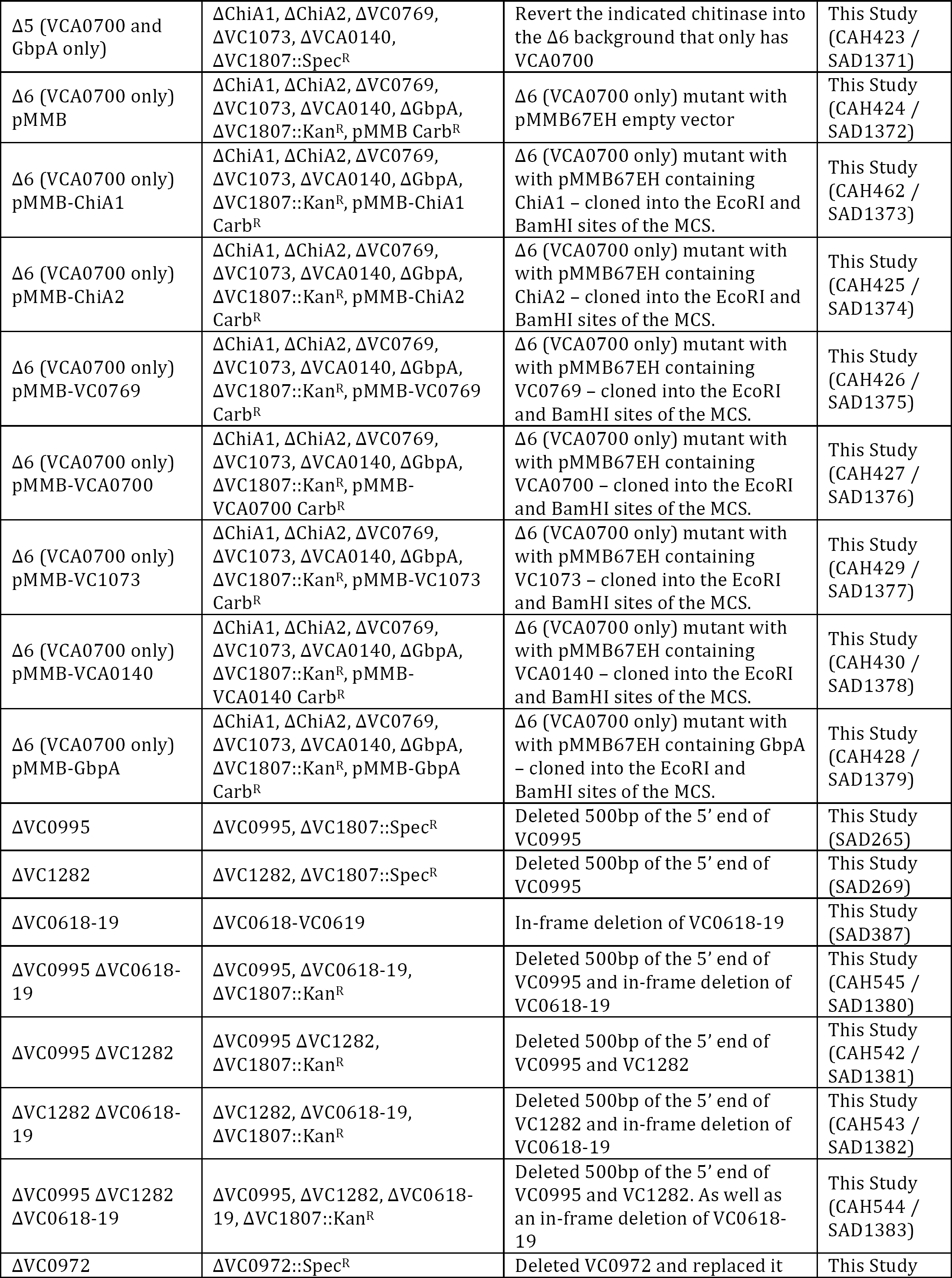
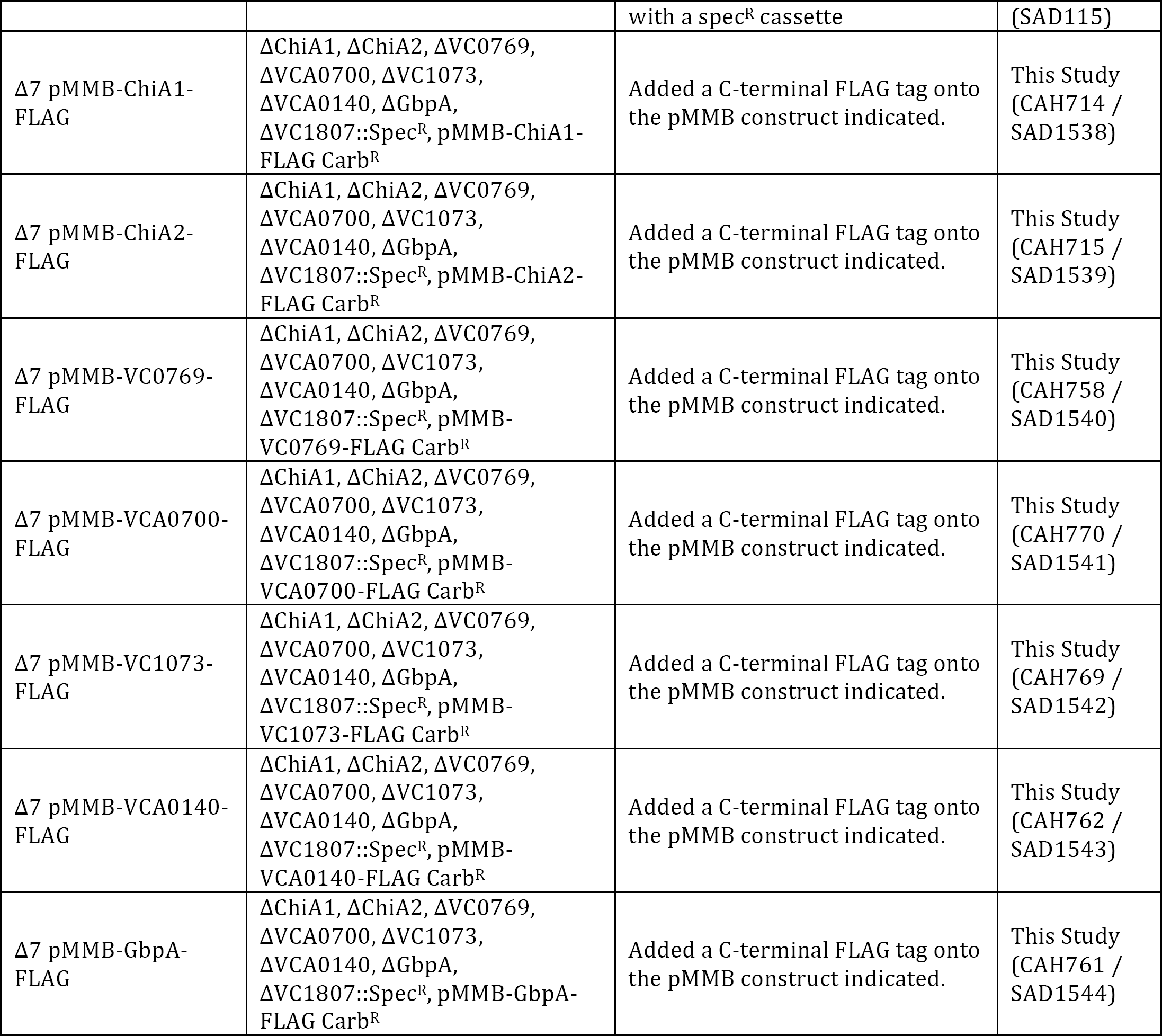
Strains used in this study

**Table S2.**
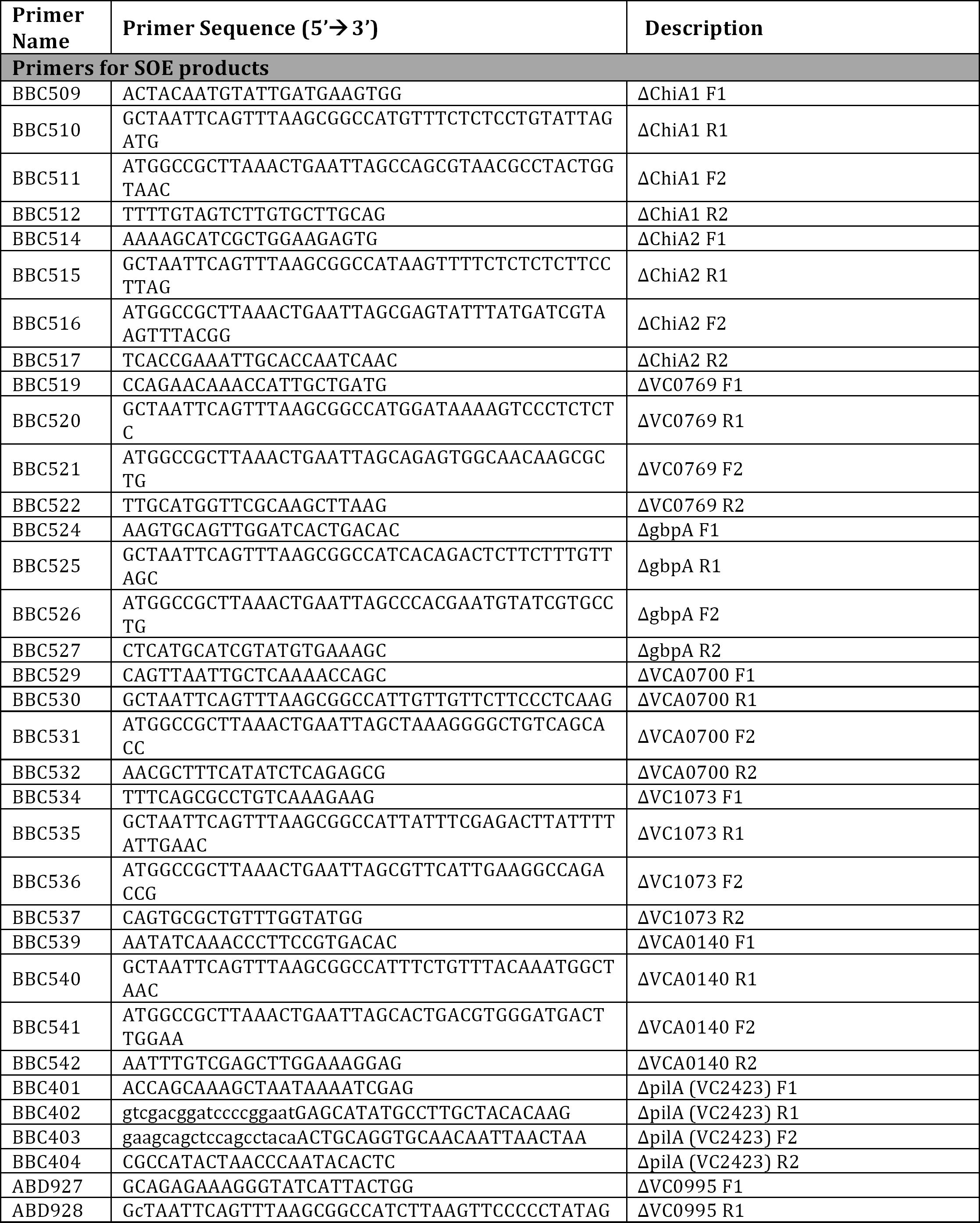
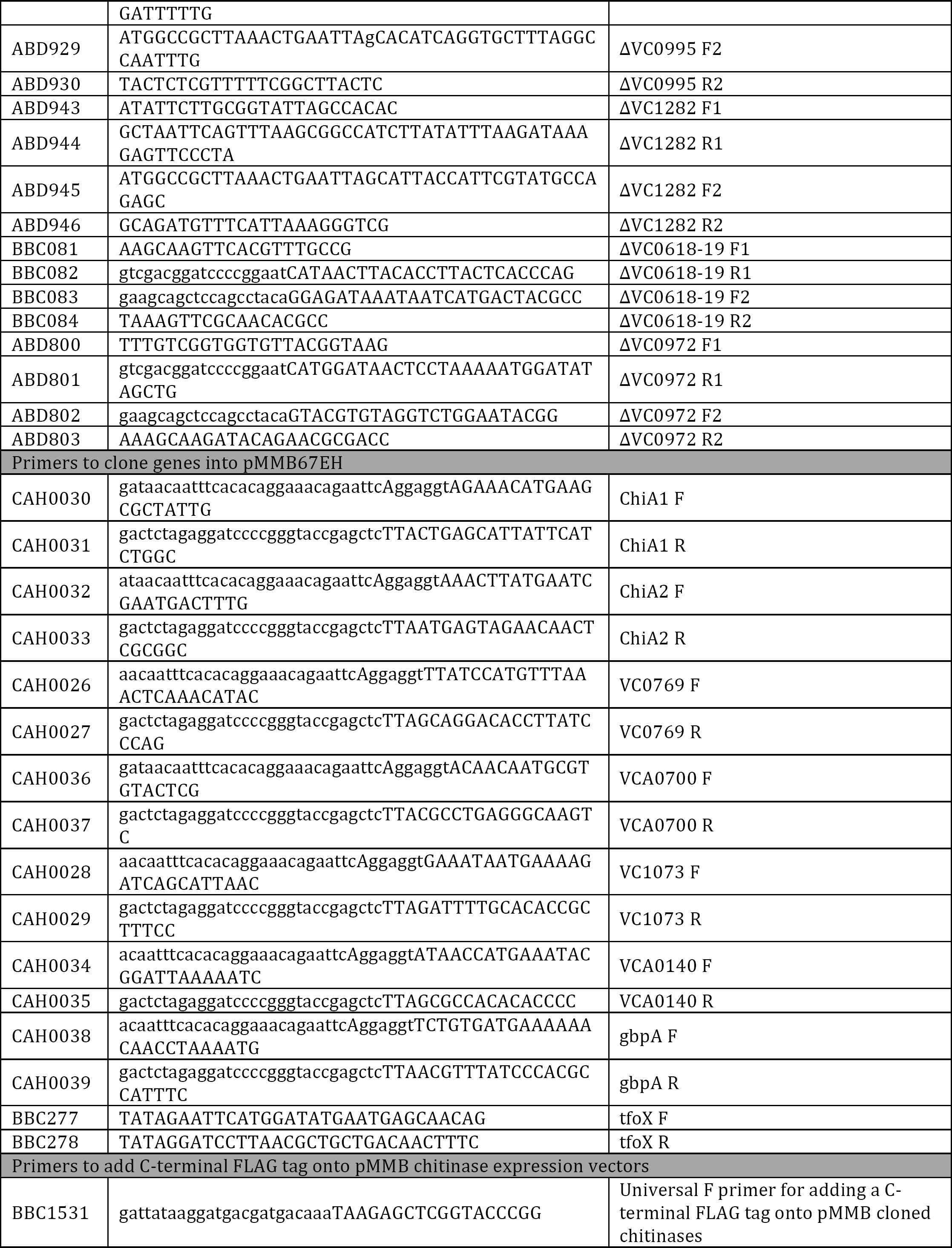
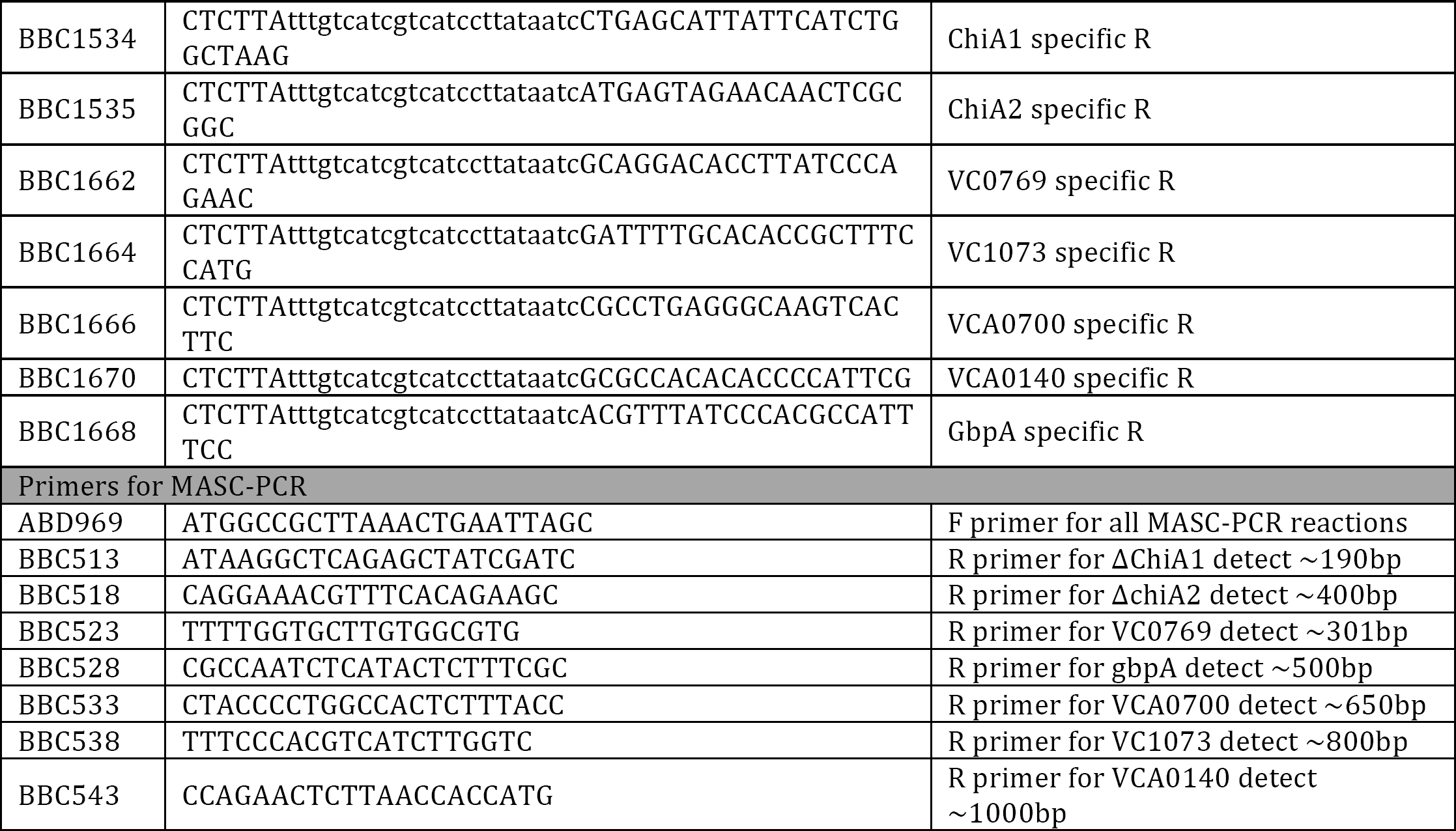
Primers used in this study

